## Extended Data Figures for "scooby: Modeling multi-modal genomic profiles from DNA sequence at single-cell resolution"

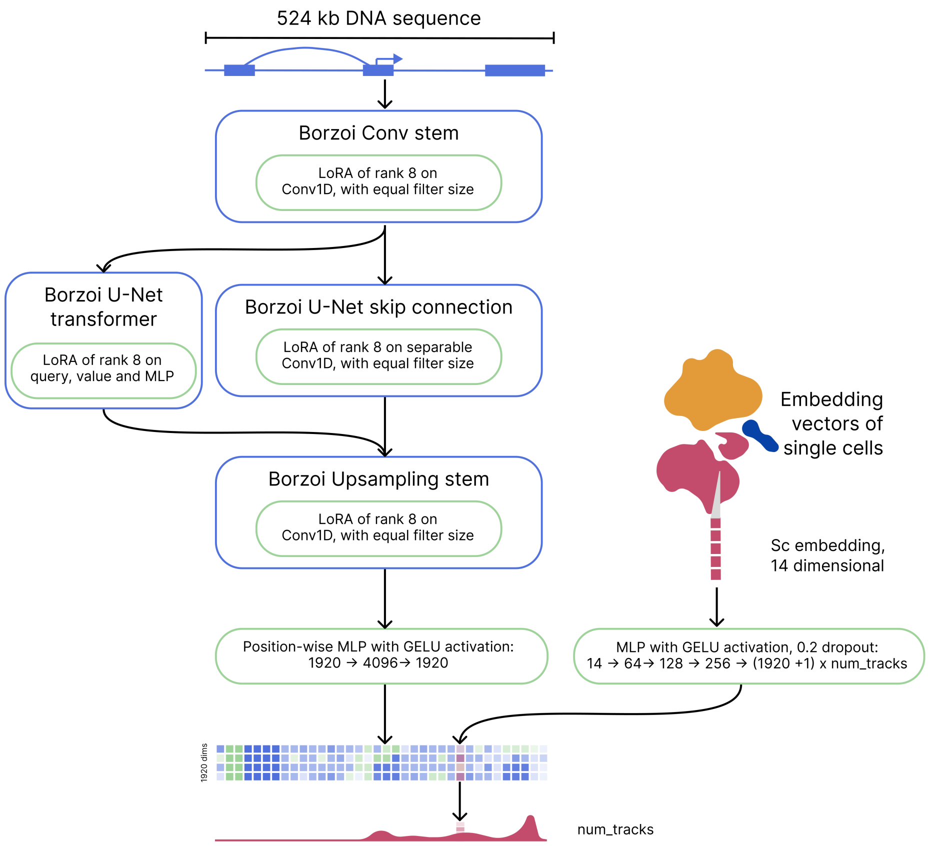


**Extended Data Fig. 1: scooby architecture overview.** Neural network diagram. Green boxes denote trainable parameters, blue boxes depict frozen (non-trained) model parts. 524kb of DNA sequence is processed by a LoRA augmented Borzoi stem with an additional MLP on the Borzoi embedding (left), whereas the single cell embedding is passed through a MLP (right) to predict the filter weights (red boxes) used to decode the sequence embedding.

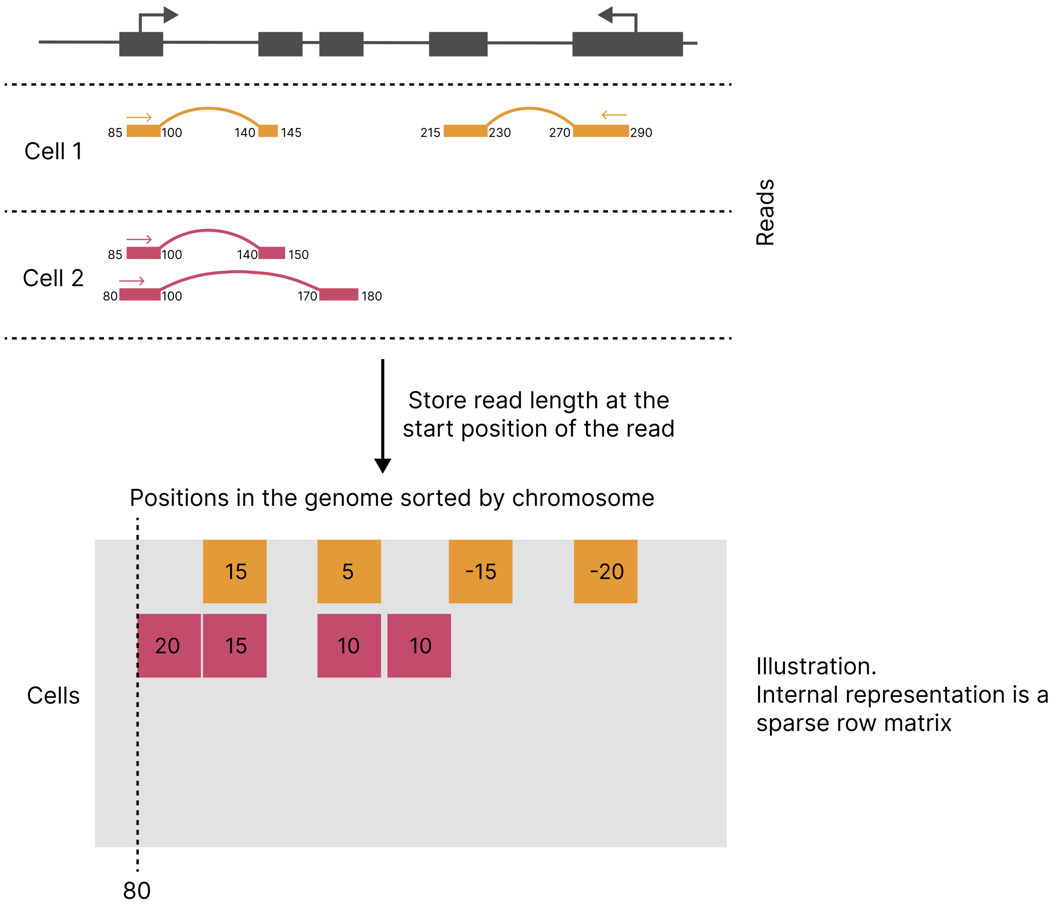


**Extended Data Fig. 2:** Memory-efficient storage of single-cell RNA-seq data using a modified snapATAC2.0 format. Each row in the resulting sparse matrix represents a cell, and each column represents a genomic position. Reads are stored at their start positions with values indicating read length. Split reads are stored as multiple entries. Negative values indicate reads mapped to the negative strand.

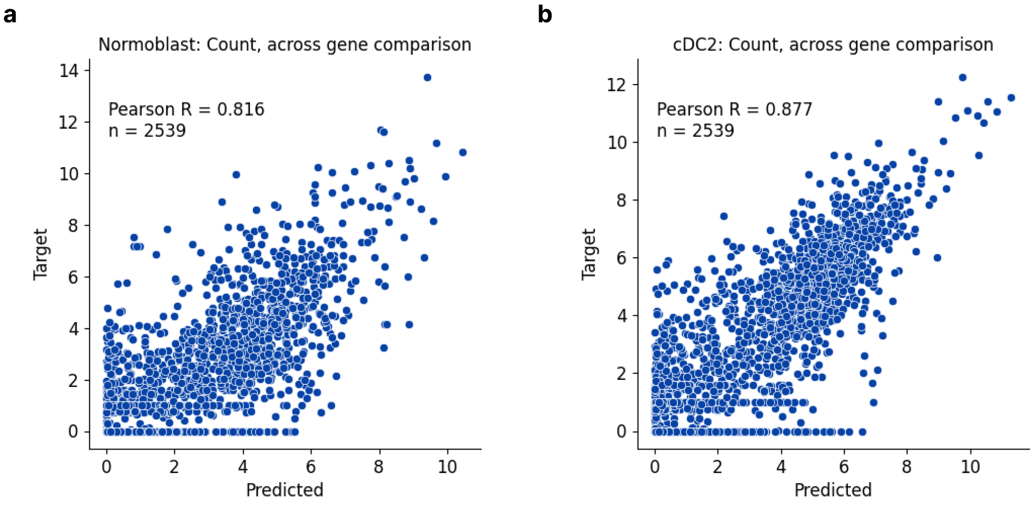


**Extended Data Fig. 3:** Log-transformed predicted versus observed counts of scRNA-seq reads overlapping exons for the worst (**a**) and best (**b**) predicted cell type.

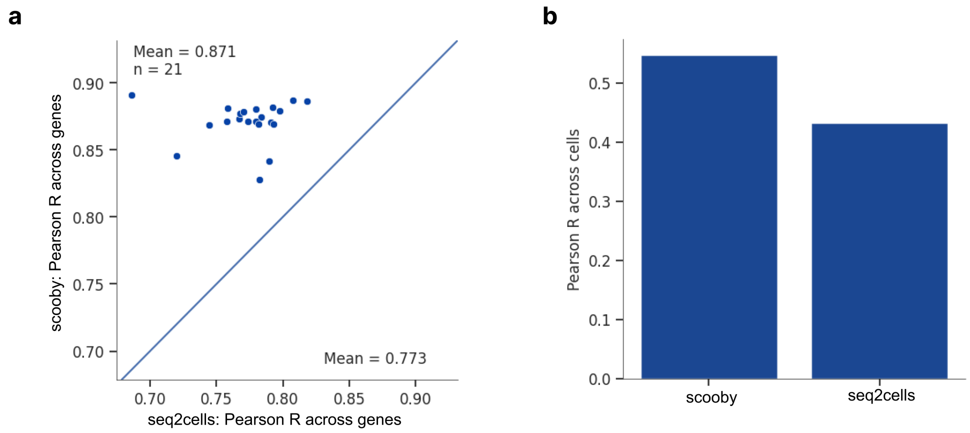


**Extended Data Fig. 4: a,** Across-gene Pearson correlation for all cell types comparing scooby and seq2cells. **b,** Between-cell-type Pearson correlation after subtracting gene and cell mean gene expression comparing scooby and seq2cells.

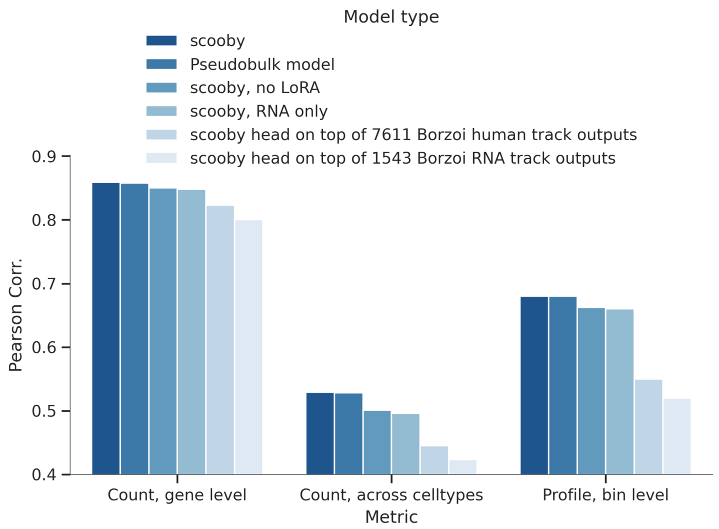


**Extended Data Fig. 5:** Performance comparison of alternative modeling approaches at predicting gene expression counts and binned profiles.

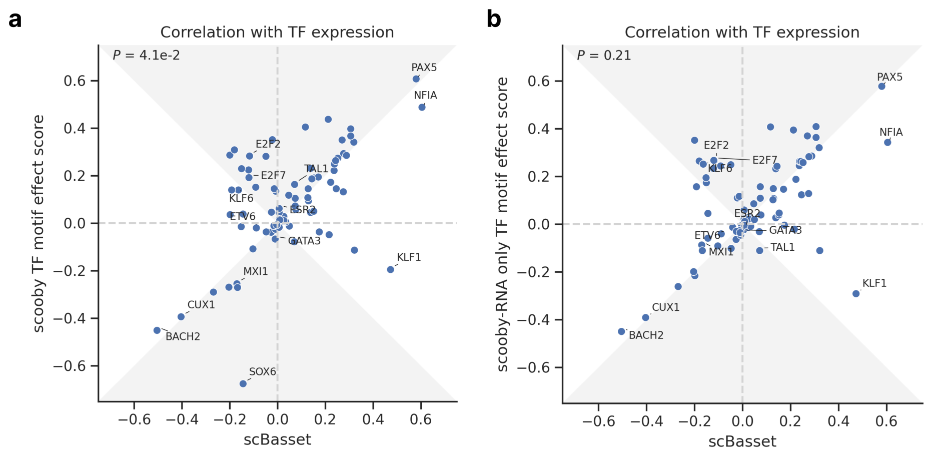


**Extended Data Fig. 6: a,** Pearson correlation of TF motif effect score with TF expression for scooby against scBasset. The gray area marks the zone of improvement. **b,** Same as **a** for a scooby trained on scRNA-seq only.

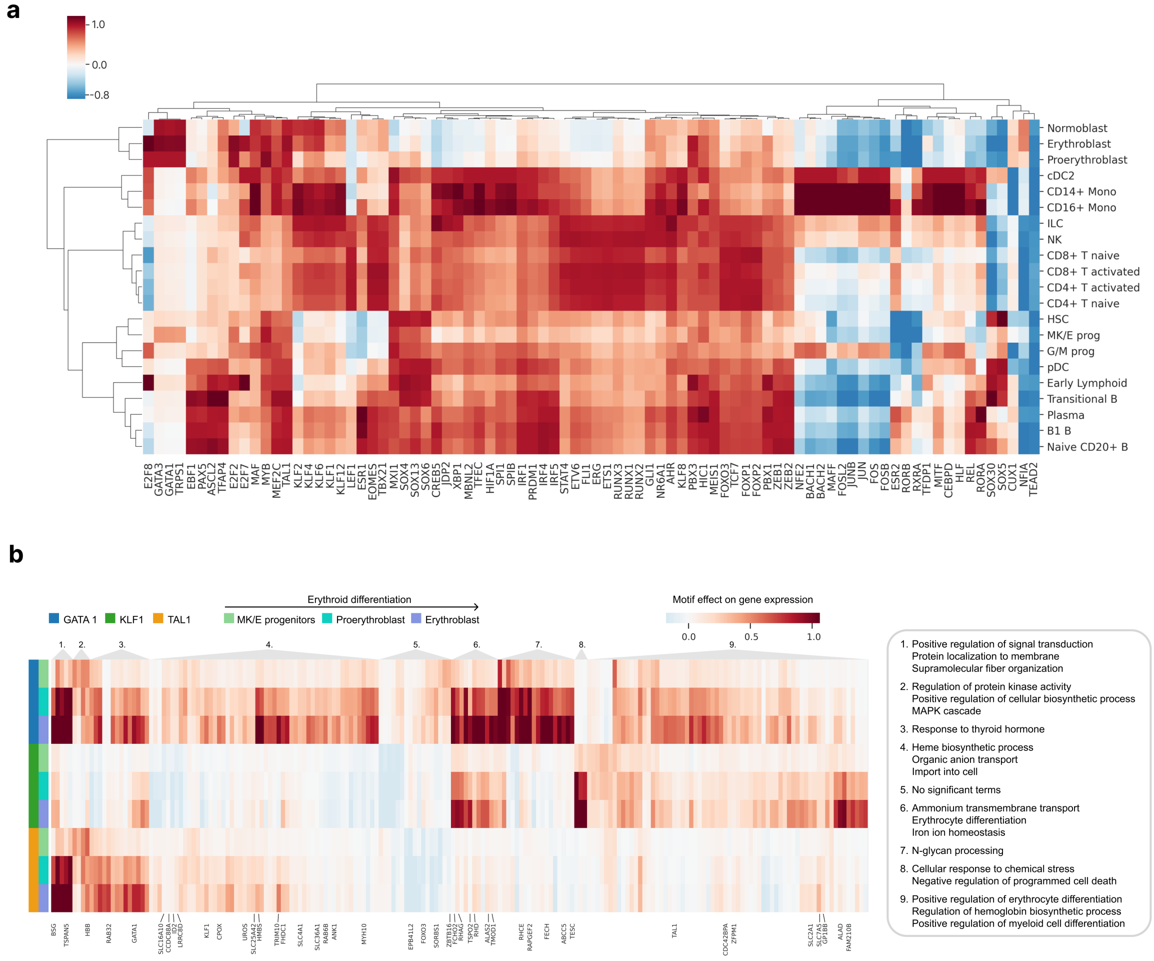


**Extended Data Fig. 7: a,** Heatmap of TF motif effect scores for differentially expressed TFs for all cell types. Genes and cell types are clustered according to their TF motif effect scores. **b,** Heatmap of genes with high motif mutation effects of GATA1, KLF1 and TAL1 in the erythrocyte lineage. Genes are clustered according to their motif effect score. The three most significantly enriched GO terms for each cluster are shown. Gene names are only shown for genes ascribed to these terms.

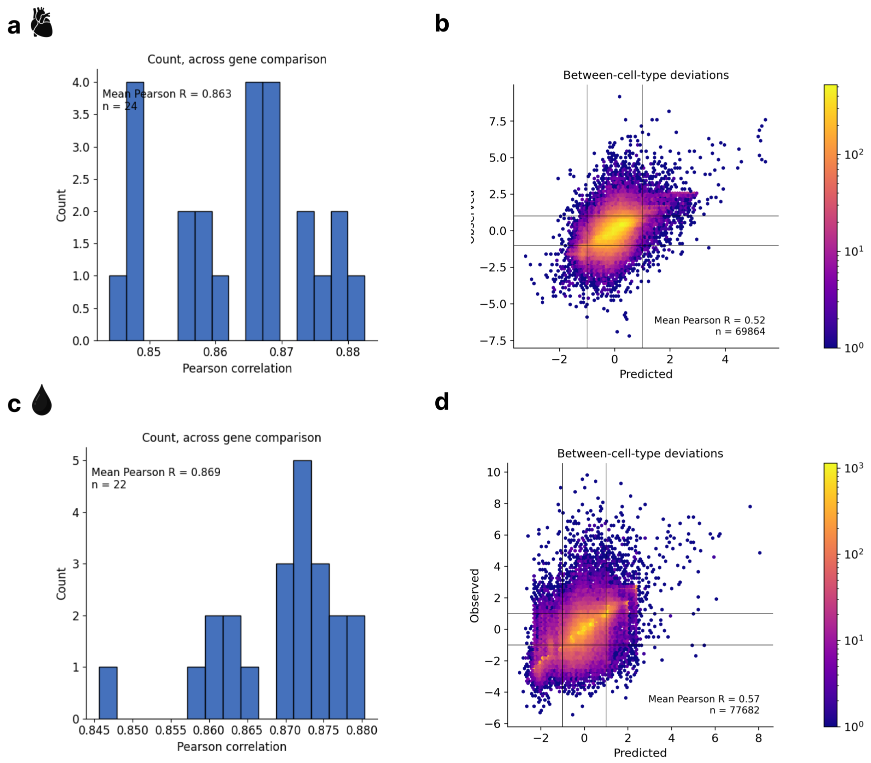


**Extended Data Fig. 8:** Distribution of gene-level Pearson correlation between log-transformed predicted and observed counts of scRNA-seq reads overlapping exons across cell types for the Epicardioids dataset (**a**) and the OneK1K dataset (**c**). Predicted against measured between-cell-type deviations of gene expression for the Epicardioids dataset (**b**) and the OneK1K dataset (**d**).

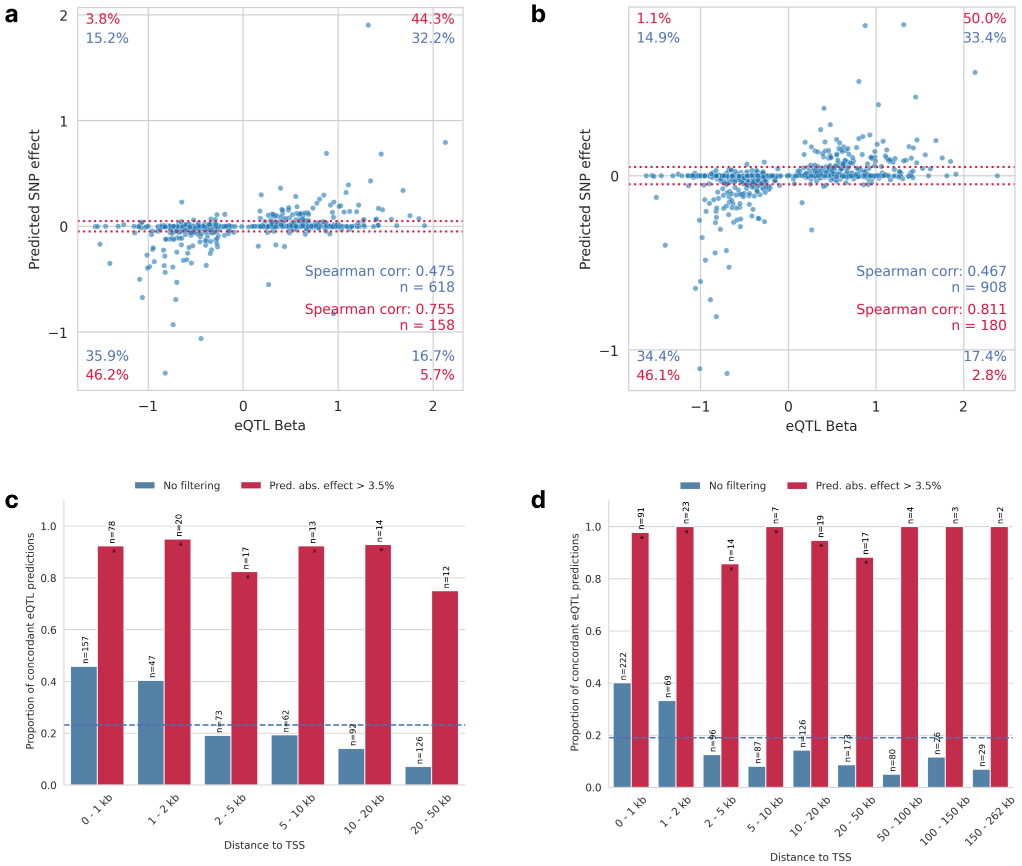


**Extended Data Fig. 9: a,** Seq2cells predicted aggregated effects (log-fold change) vs. observed whole-blood eQTL effect sizes. Red dotted lines mark thresholds below which predicted fold-changes are deemed negligible (absolute fold change 3.5%; matching the threshold by Schwessinger et al. for comparability). Percentages quantify variants within each quadrant: blue - all variants; red - variants passing the 3.5% predicted effect threshold. **b,** Same as **a,** but for Borzoi. **c,** Proportion of concordant seq2cells eQTL predictions (same direction as observed), as a function of distance to the transcription start site when filtering for non-negligible predicted effect (red) or without filtering (blue). Dashed blue line indicates the mean proportion of concordant eQTL predictions across all distances (0.23). Stars indicate significance over random performance (Binomial test). **d,** Same as in **c,** but for Borzoi.

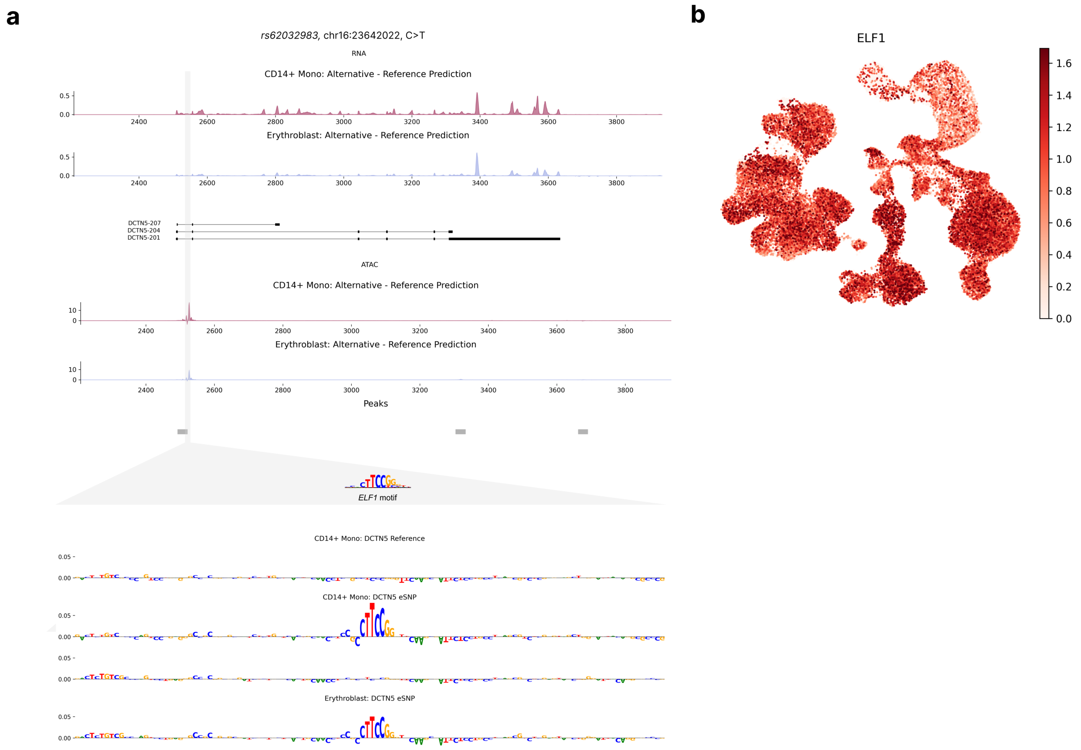


**Extended Data Fig. 10: a**, Predicted fold change in gene expression (top) and accessibility (bottom) between the alternative and reference alleles of variant rs62032983 in CD14+ Monocytes and Erythroblasts. Sequence attributions revealed the destruction of an *ELF1* motif to affect model outputs across cell types (Methods). **b**, UMAP of observed normalized *ELF1* expression levels.

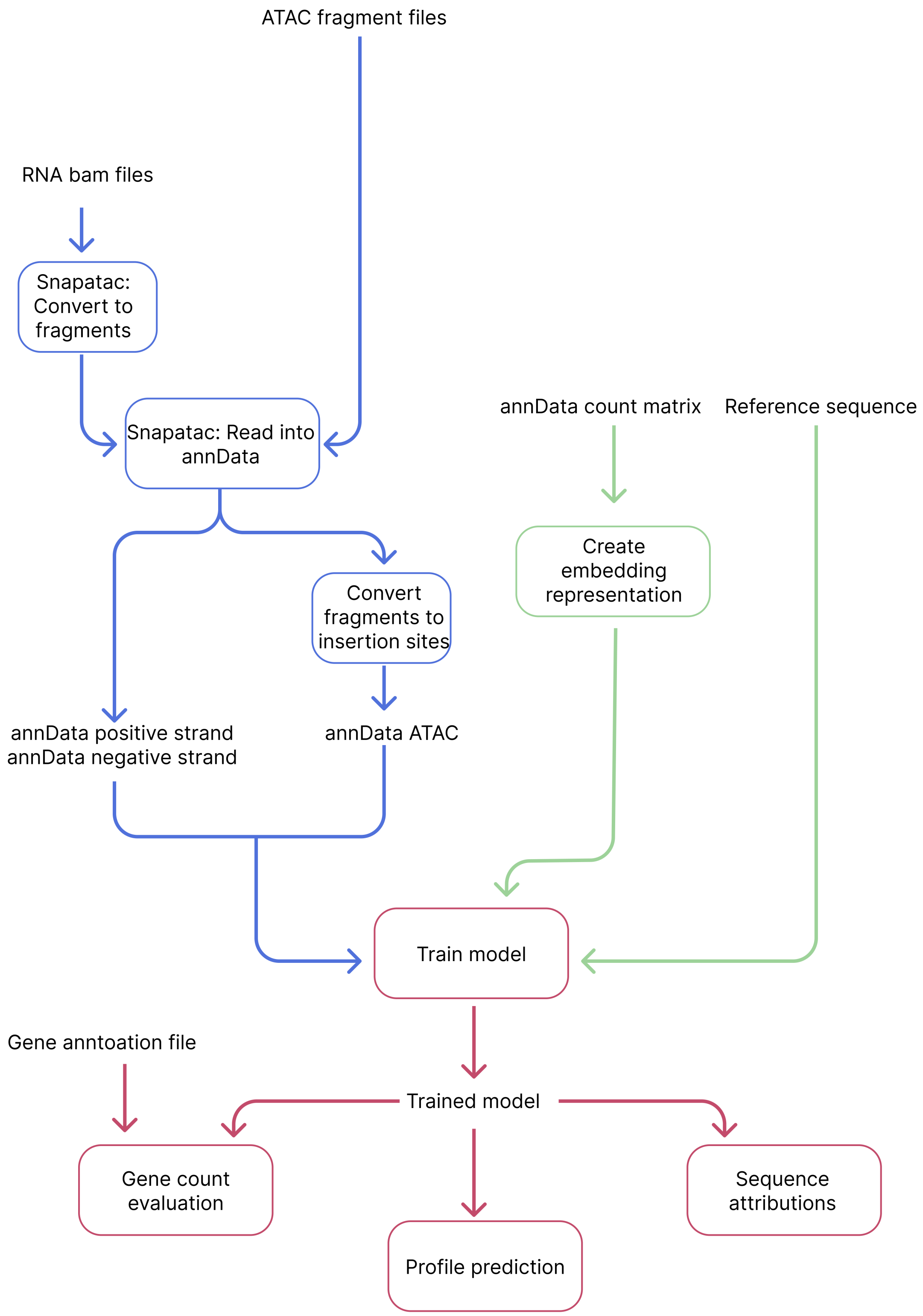


**Supplementary Fig. 1: scooby framework flowchart.** Diagram illustrates the workflow for generating cell embeddings and preparing RNA-seq and ATAC-seq data for training, including filtering, embedding generation, and coverage extraction.
